# A General-Purpose Non-invasive Neurotechnology Research Platform

**DOI:** 10.1101/2024.01.01.573494

**Authors:** Gerwin Schalk, Shiyun Shao, Kewei Xiao, Jintao Li, Jiaxin Xie, Yinkui Guan, Zehan Wu, Liang Chen, Xingyi Zhong, Ce Xu, Guangye Li, Huan Yu

**Affiliations:** Chen Frontier Lab for Applied Neurotechnology, Tianqiao and Chrissy Chen Institute, Shanghai, China; Department of Neurosurgery, Huashan Hospital, Fudan University, Shanghai, China; University of Electronic Science and Technology of China, Chengdu, China; Institute of Robotics, Shanghai Jiao Tong University, Shanghai, China; Sleep and Wake Disorders Center of Fudan University, Shanghai, China; Department of Neurology, Huashan Hospital, Fudan University, Shanghai, China; National Center for Neurological Disorders & National Clinical Research Center for Aging and Medicine, Huashan Hospital, Fudan University, Shanghai, China

## Abstract

This article describes our initial work toward a general-purpose platform for non-invasive neurotechnology research. This platform consists of a multi-modal wireless recording device, an associated software API, and full integration into BCI2000 software. The device is placed on the forehead and features two electroencephalographic (EEG) sensors, an inertial movement sensor (IMU), a photoplethysmogram (PPG) sensor, a microphone, and vibration-based feedback. Herein, we demonstrate different technical characteristics of our platform and its use in the context of sleep monitoring/modulation, simultaneous and synchronized recordings from different hardware, and evoked potentials. With further development and widespread dissemination, our platform could become an important tool for research into new non-invasive neurotechnology protocols in humans.

## 1. Introduction

Recordings from the surface of the scalp (electroencephalography (EEG)) have been used for decades in the clinic and in research laboratories. For example, clinicians commonly interpret EEG recordings and other physiological signals to generate useful diagnostic information about specific sleep disorders (Rundo and Downey III, 2019). Likewise, thousands of research studies have shown that the EEG holds substantial information about a person’s function or dysfunction, such as parameters of depression (deAguiar Neto and Rosa, 2019) or cognition (Doan et al., 2021; Jeong, 2004). These studies have also shown that EEG can be used, together with appropriate conditioning protocols, to replace, restore, improve, enhance, or supplement functions lost due to different neurological disorders (Wolpaw and Wolpaw, 2012), such as to restore motor or speech function that is lost or impaired after stroke (Bundy et al. (2017) or Musso et al. (2022), respectively).

In summary, there is now ample if not overwhelming evidence to suggest that EEG could support functions that, in principle, could prove useful not only in the context of (a relatively limited number of) research studies or clinical evaluations, but could also improve the lives of a large number of people in their home. However, it is currently largely unclear how to transfer the potential benefits suggested or even realized by ongoing clinical practice or by scientific experimentation to practical in-home solutions. Determining how to generate such solutions requires, for each use case, large-scale evaluations of different neuroscientific protocols and engineering approaches in people’s homes.

Traditional research-focused EEG systems can support a wide range of research, but they are too impractical for large-scale use outside the laboratory (see Fig. 1). To address this issue, over the past 20 years, an increasing number of manufacturers have produced consumer-centric hardware that is meant to support EEG-based applications in people’s homes and/or other natural environments. These consumer EEG systems generally fall into two categories.

**Figure 1.**
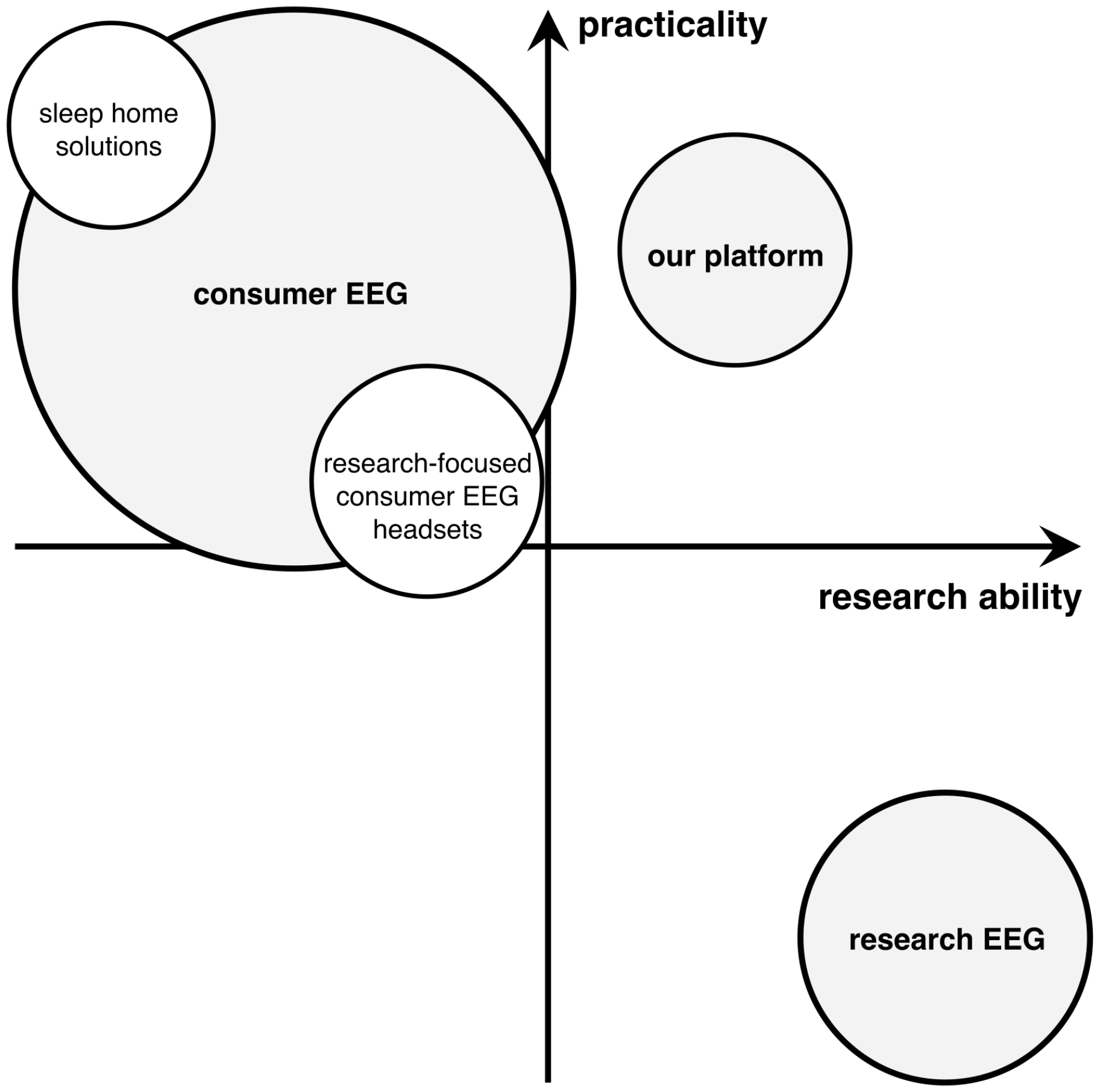
Existing research and consumer EEG solutions, and our platform. Traditional research EEG systems support a wide gamut of research, but they are relatively impractical to use. Existing consumer EEG solutions (sleep home solutions as well as other consumer EEG headsets) are more practical, but have limitations in the research they can support. Our hardware/software platform is designed for a balance of research ability and practicality.

The first category of consumer EEG systems is sleep home solutions that are meant to complement the more traditional polysomnographic (PSG) evaluations that are performed at sleep labs. For example, companies including Compumedics (Compumedics, 2023), Dreem/Beacon Signals (Arnal et al., 2020; Beacon Biosignals, 2023), Shenzhen EEGSmart Technology (Shenzhen EEGSmart Technology, 2019), FlectoThink (FlectoThink, 2022), and VentMed (Hunan VentMed Medical Technology, 2017) developed forehead patches or headbands for in-home sleep monitoring. These commercial in-home sleep solutions are usually quite practical, but they often have limitations in signal quality and/or their ability to use them for research purposes. For example, most of them do not have an API that provides real-time access to raw signals. There are also research laboratories that have been focusing on different solutions for similar specific purposes (da Silva Souto et al., 2021; Kwon et al., 2023; Nakamura et al., 2017; Tabar et al., 2023; Xu et al., 2023), but these academic solutions do not support a wide array of applications, and are typically not available to others.

The second category of consumer EEG systems is commercial devices that evolved from the context of laboratory EEG research. These devices include systems from different manufacturers such as Emotiv (EMOTIV Inc., 2023), g.tec (g.tec, 2023), Muse (InteraXon Inc., 2023), Neurosky (NeuroSky Inc., 2023), OpenBCI (OpenBCI, 2022), and Wearable Sensing (Wearable Sensing, 2023). These devices are typically better suited for research (e.g., they all have an API that supports real-time access to signals), but they are usually less practical than sleep home solutions since they have wet electrodes or electrodes that are placed in the hair. An interesting exception is the recent introduction of the EEG-enabled headphones by Neurable (Neurable, 2023), but more information about the device’s capabilities and limitations is needed.

None of these consumer EEG solutions are currently packaged with powerful general-purpose closed-loop software that facilitates the wide array of evaluations necessary for development of practical neurotechnology solutions. In summary, there is currently no solution that can simultaneously provide full research/clinical abilities and clinical-grade practicality/robustness. We are painfully aware that it will likely be impossible to fully address both of these requirements simultaneously. For example, comprehensive polysomnography (PSG) evaluations require measurements of chest movements (e.g., to disambiguate central and obstructive sleep apnea) or of leg movements (e.g., to provide evidence for Restless Leg Syndrome). It is almost certain that such (and other) measurements cannot be accomplished using sensors that are placed in a completely different location such as on the head. Likewise, optimization of practicality almost certainly requires placing electrodes outside the hair (e.g., on the forehead or around/inside the ear), but these locations will not provide optimal access to EEG signals that are typically detected in more central locations (Schalk et al., 2023).

Within these general constraints, with the work described in this paper, we set out to develop a platform that combines multi-modal clinical-grade signal acquisition, high practicality and robustness, and significant research abilities. This platform consists of a forehead patch that provides access to multi-modal signals, a software API, and powerful closed-loop software that is optimized for neurotechnology research. We expect that our system will greatly facilitate research into new approaches for home sleep monitoring and modulation, as well as other home-based diagnostic and treatment solutions. Thus, we hope that with further continuation of the work described herein, we will be able to hasten the transition of laboratory findings and clinical approaches into clinically and commercially successful solutions that will improve the lives of many people.

## 2. Methods

### 2.1. System

#### 2.1.1. Hardware

To support clinical-grade and practical data collection, we developed multi-modal recording hardware that is capable of efficient closed-loop operation. It is relatively small (89 mm * 47 mm * 5 mm) and light-weight (30.6 grams) and, together with a disposable electrode patch, is being placed on the forehead (see Fig. 2). It is powered by a rechargeable battery that lasts more than 10 hours, and it communicates to a host using Bluetooth 5.0 BLE via a Bluetooth dongle (PCs) or directly (iOS/Android). The device’s electronics are placed on a flexible printed circuit board (PCB) and are enclosed in medical-grade flexible silicone (Shore hardness of 80A) to ensure the flexibility and comfort of the equipment and to accommodate the different shape of people’s forehead.

**Figure 2.**
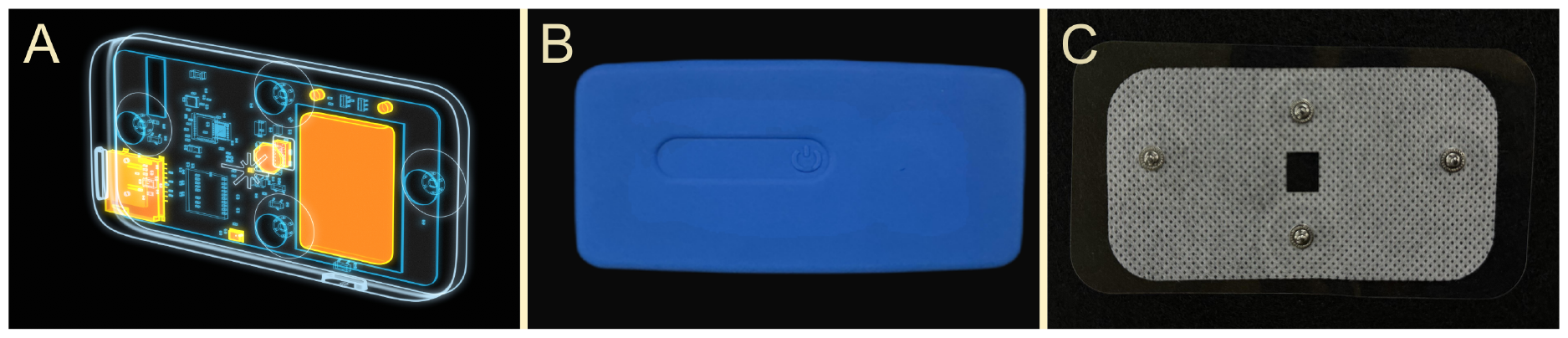
Multi-modal recording hardware. A: X-Ray diagram of the device and its components. B: Front of the device. C: Front of the electrode patch that clips to the back of the device.

Our device has several sensors that support EEG recordings, photoplethysmography (PPG), detection of movements (using an inertial measurement unit (IMU)) and sounds (using a microphone), and it can provide vibration feedback using a linear motor.

The EEG signals are detected from Fp1 and Fp2 (referenced to Fpz) using a 24-bit ADC (ADS1299-4, Texas Instruments, USA), and are sampled at 500 Hz after low-pass filtering at 100 Hz. The input range of the EEG signal is *±*600*mV*_*pp*_, and the input voltage noise (RTI) noise is *<* 1*μV*_*pp*_.

PPG recordings are supported by a 3-wavelength optical sensor (Max30101, Analog Devices, USA) that samples the degree of reflection to light emitted at 537 nm (green light), 660 nm (red light), and 880 nm (infrared light) wavelengths, respectively, at 100 Hz. Together with appropriate algorithms, PPG recordings can be used to derive heart rate and heart rate variability (Biswas et al., 2019; Pankaj et al., 2022; Ye et al., 2016), as well as blood oxygenation (SpO2) (Alkhoury et al., 2021; de Kock and Tarassenko, 1991; Wukitsch et al., 1988).

IMU recordings are accomplished using an integrated 9-axis motion tracking device (ICM-20948, TDK InvenSense, Japan) with a sampling rate of 50 Hz. Using appropriate methods such as non-linear complementary filters (Mahony et al., 2008) or gradient-descent orientation filters (Madgwick, 2010), these IMU signals can be converted into absolute position and movement of the device (and thus, the user’s head).

The sound signal is detected using a MEMS microphone (MP23ABS, STMicroelectronics, Italy) with a sampling rate of 1000 Hz. The sound signal can be used to detect ambient noise and, using appropriate algorithms, the user’s snoring.

The device can provide vibration feedback using a linear motor that supports three modes of vibration (constant vibration, pulse vibration, and sinusoidal vibration).

A single-use electrode patch provides a robust interface with the user’s forehead. It consists of four electrodes (Fp1, Fp2, Fpz (reference) and ground), medical non-woven fabric, and a medical adhesive layer. Electrodes are made of Ag/AgCl and are in contact with the scalp via a solid PVA-H hydrogel.

#### 2.1.2. Hardware Testing

In initial evaluations, we determined three EEG measurement characteristics, namely amplitude accuracy, noise root-mean-square (RMS), and common-mode rejection ratio (CMRR), and we compared our device’s characteristics to those of three traditional research-focused EEG acquisition devices: 1) D440, Digitimer Ltd., UK; 2) g.HIamp, g.tec, Austria; and 3) NeuSen W, Neuracle, China.

To measure amplitude accuracy, we used a signal generator to produce a 100 *μV* 10 Hz square wave signal *U*_*in*_. We connected the generator’s signal pin to one recording channel of our device, and the generator’s reference signal pin to our device’s reference and ground channel. We then measured the amplitude of this signal as *U*_*m*_, and determined the maximum deviation from the expected value to calculate the maximum error *δ*_*m*_:

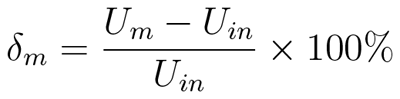

Finally, we calculated amplitude accuracy using 100 *− δ*_*m*_.

To measure noise RMS, we shorted out our device’s signal input channel, reference channel, and ground, and then calculated RMS amplitude *N*_*rms*_ given the following equation:

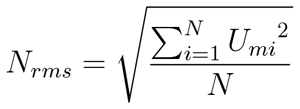

Finally, to measure CMRR, we used a signal generator to produce a sinusoidal signal (3 Volt peak-to-peak, 50 Hz), and connected the generator’s signal channel to our device’s signal and reference channels, and the signal generator’s reference channel to our device’s ground. We then used our device to measure the peak-to-peak voltage of this signal as *U*_*c*_. We then changed the signal’s amplitude to 3 mV, and used our device to measure the peak-to-peak voltage of this signal as *U*_0_. Finally, we calculated CMRR using the following equation (*K* = 1000):

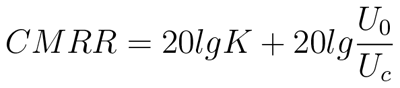

We also determined timing characteristics that are important for potential closed-loop application of our system. To do this, we tested the average duration of each transmitted (20 ms long) data block, its jitter/standard deviation, as well as the fraction of dropped blocks. We performed this testing when the device and associated Bluetooth receiver were right next to each other or 2 meters apart. The results are shown in Table 1.

**Table 1.** Timing characteristics and dropped packets for device operation that is close to or 2 meters away from the Bluetooth receiver.

|  | close | 2 meters |
| --- | --- | --- |
| mean | 20.00 ms | 20.02 ms |
| jitter/standard deviation | 0.70 ms | 4.22 ms |
| packets dropped | 0.0% | 0.6% |

They demonstrate that, even though the device communicates through a wireless link, its timing characteristics are excellent and there is no or only minimal packet loss when the device is within a reasonable range to the receiver.

#### 2.1.3. Software

*API* To support the hardware functions of our recording device, we implemented a C API. This API can identify all currently available devices, connect to a specific device, start data streaming, operate the vibration motor, and perform other functions. The specific functions included in our API are:

- `Open()` This function opens the serial port provided by the Bluetooth dongle and sets necessary parameters for serial communication such as baud rate.
- `FindDevice()` This function finds all devices that are powered on and are within Bluetooth range, and determines whether the device we want to connect to exists based on the device’s MAC address.
- `Connect()` This function connects to the desired device.
- `CheckConnect()` This function checks whether the device and the Bluetooth receiver are properly connected.
- `Start()` This function turns on data collection and transmission.
- `End()` This function stops data collection and transmission.
- `TurnOnMotor()` This function turns on the vibrating motor.
- `TurnOffMotor()` This function turns off the vibrating motor.
- `Disconnect()` This function disconnects a device.
- `Close()` This function closes the serial port.

#### 2.1.4. Closed-Loop Software

To enable comprehensive closed-loop neurotechnology capabilities, we developed full support for our hardware device in BCI2000. BCI2000 is a general-purpose software platform for closed-loop neurotechnology (Schalk et al., 2004; Schalk and Mellinger, 2010), and has been in active development for close to 25 years. Over this period, BCI2000 has supported experiments reported in more than 1000 peer-reviewed publications (Brunner and Schalk, 2018), including many highly influential studies in the neurotechnology literature (e.g., Herff et al. (2015); Leuthardt et al. (2004); Miller et al. (2010); Wolpaw and McFarland (2004)).

With appropriate hardware, BCI2000 can acquire signals from the brain, body physiology, or behavior, process them in meaningful ways, and use the outputs to control the timing or other properties of feedback. These capabilities are highly flexible and performant, and execute robustly even with demanding requirements. The comprehensive integration of our hardware with BCI2000 provides many useful functions. For example, it can:

- Acquire and synchronize all signals provided by our device, i.e., two channels of EEG, accelerometer/gyroscope/magnetometer (providing absolute body position), IR/red/green light (photoplethysmograph (PPG), providing heart rate, heart rate variability, and SpO2), and microphone
- Provide highly customized tactile feedback through the vibration motor
- Synchronize signals from our device with behavioral measurements acquired from many supported devices such as eye trackers, data gloves, or wearable movement sensors
- Calculate spectral amplitude/power/phase using different algorithms (e.g., bandpass-filtering and Hilbert transform, FFT, or AR spectral estimation)
- Compute different types of statistics of these measurements
- Provide auditory, visual, or other stimulation contingent on the results of these statistics

Users can harness these capabilities without delving into programming intricacies, or can enhance them using documented interfaces in C++, Python, Matlab, and Simulink. The versatility of the filtering tools extends to both real-time brain signal data and offline data analysis, offering a streamlined avenue for algorithm optimizations. Moreover, the inclusion of comprehensive scripting features empowers users to craft sophisticated, fully automated experimental protocols.

BCI2000 excels in demanding experimental scenarios, offering exceptional performance that ensures swift feedback with minimal latencies and jitter (Wilson et al., 2010). For example, under optimized settings, the audio output jitter remains below 1 ms, while stimulation latency is kept under 3 ms. Notably, BCI2000 includes a timing certification system that can assess the timing of any BCI2000 configuration, encompassing both hardware and software components.

## 3. Results

### 3.1. Sleep Monitoring and Modulation

We here demonstrate the application of our platform in three contexts. The first of these contexts is sleep monitoring and modulation.

We spend almost a third of our life sleeping. It is now clear that sleep is critical to our health and quality of life, and that poor sleep can cause reduced productivity and increased mortality (De Fazio et al., 2022; Kwon et al., 2023). Unfortunately, many people have insufficient amount of sleep, or suffer from different types of sleep disorders. Indeed, it is estimated that approximately 10% of the adult population suffer from insomnia, and an additional 20% experience occasional insomnia symptoms (Morin and Jarrin, 2022). Obstructive sleep apnea (OSA), one of the most common sleep disorders, is estimated to affect 936 million adults aged 30–69 worldwide (Benjafield et al., 2019; Surani and Taweesedt, 2022). Despite the high prevalence of these problems, 80% to 90% of people remain underdiagnosed and undertreated (Kwon et al., 2023; Senaratna et al., 2017). One of the main reasons for this important issue is the lack of easily accessible tools for evaluating different approaches to sleep monitoring and sleep modulation in people’s homes (Kwon et al., 2023, 2021).

The hardware/software platform described in this paper allowed us to begin addressing this issue by facilitating the design and clinical evaluation of our own sleep solution that supports monitoring of EEG signals, heart rate, SpO2, and head movements, and that provides auditory stimulation that seeks to enhance EEG delta activity during slow wave sleep, similar to protocols described in previous research (Garcia-Molina et al., 2018; Ngo et al., 2013; Papalambros et al., 2019; Santostasi et al., 2016). Figure 4 shows examples of results of our ongoing evaluations of these sleep monitoring and modulation functions.

**Figure 3.**
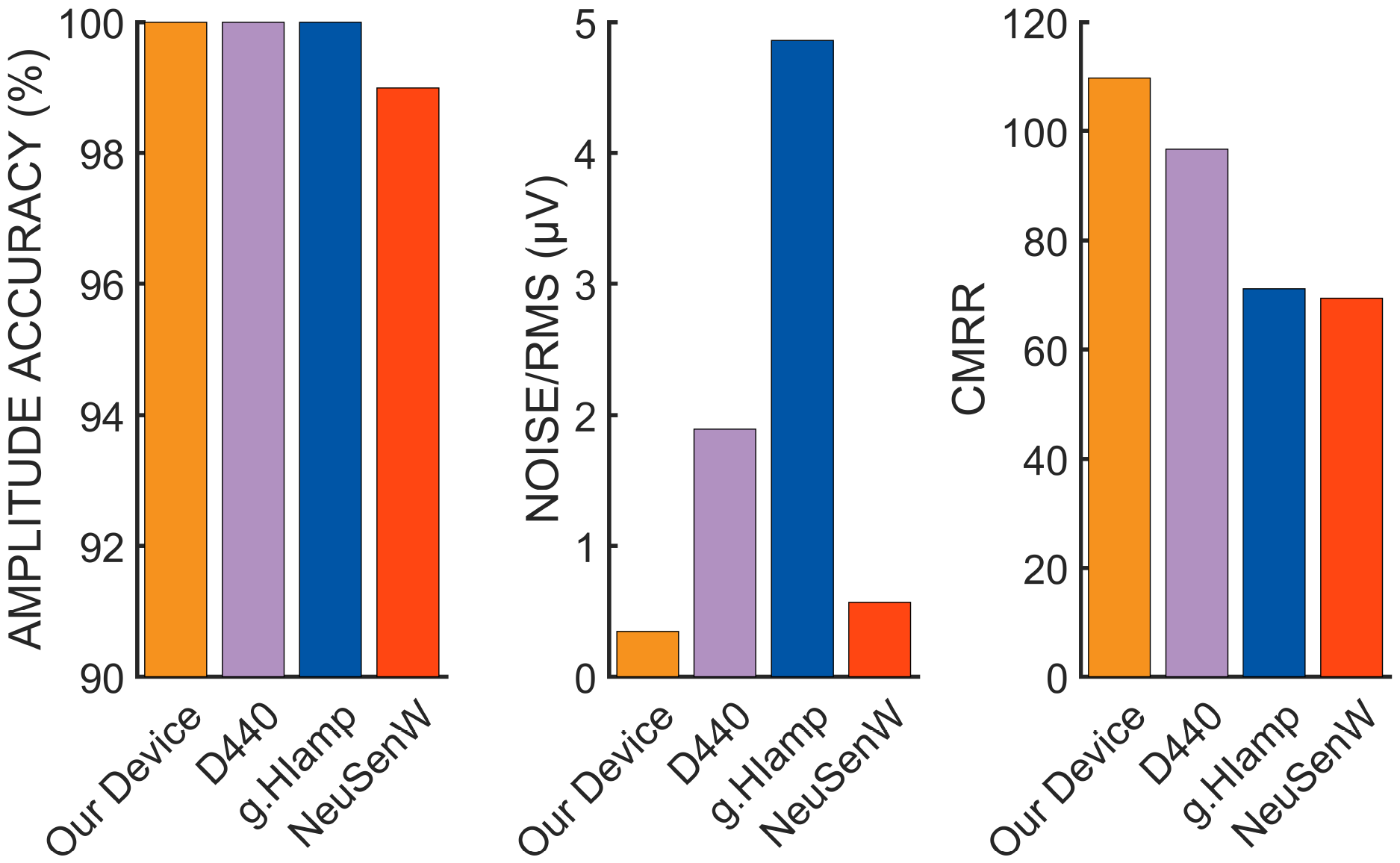
Amplitude accuracy, noise/RMS amplitude, and CMRR for our device and three research-focused devices.

**Figure 4.**
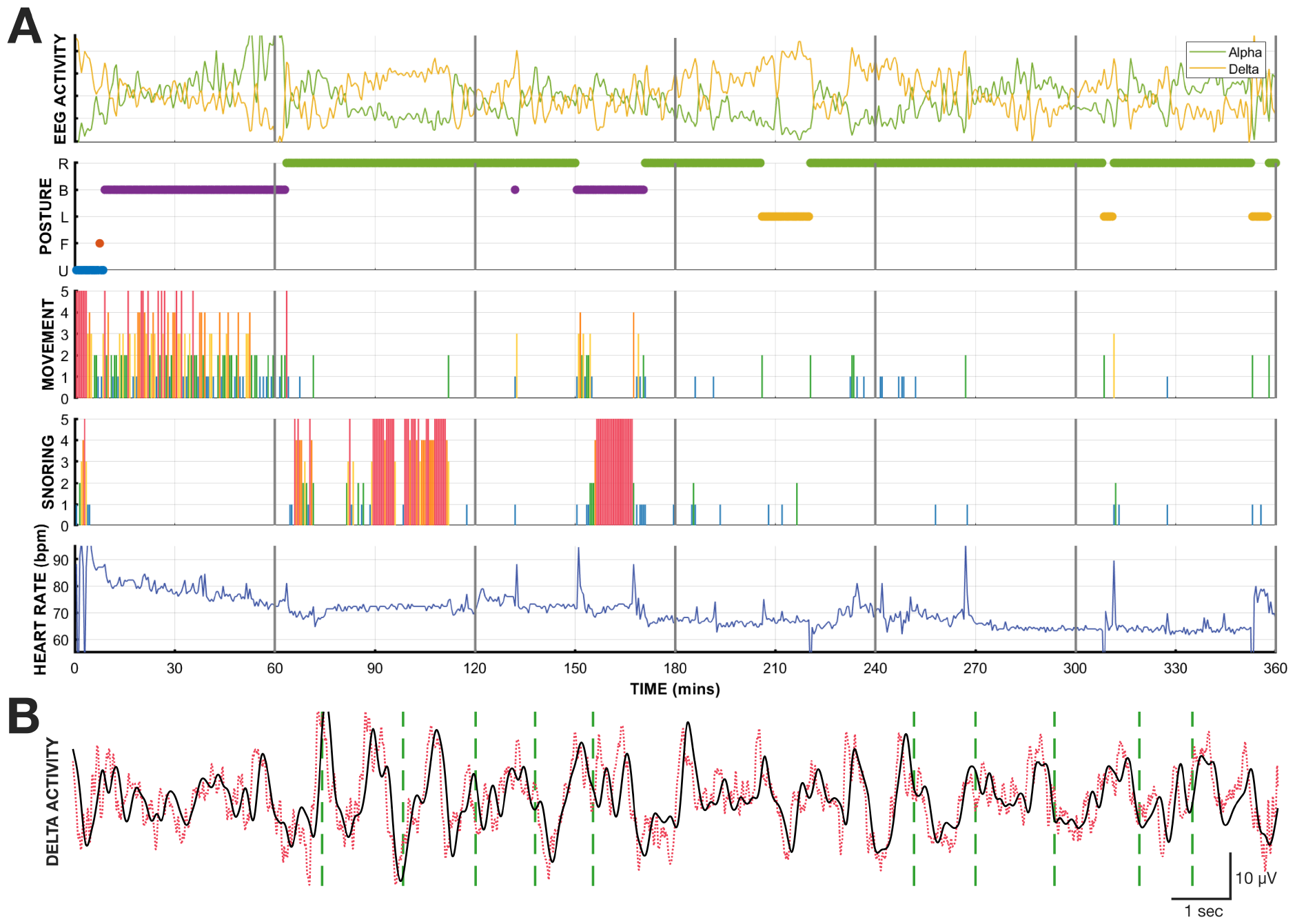
Sleep monitoring and modulation. A: Concurrent acquisition of EEG activity (filtered here in the alpha (8-12 Hz) and delta (0.5-4 Hz) ranges), posture, movement, snoring, and heart rate (top to bottom panels, respectively) in one subject during 6 hours of sleep. B: EEG signals (0.3-35 Hz bandpass, red dotted trace) and EEG in the delta range (black solid trace) in one subject. Dashed green lines indicate times of auditory stimulation at certain phases of delta activity.

### 3.2. Synchronized Acquisition of Signals From Different Devices

The second context involves the capability to acquire and synchronize signals from different devices, which enables or facilitates a number of neurotechnology applications. Specifically, our platform supports the synchronized acquisition of all EEG and physiologic/behavioral measurements provided by our device with those of other behavioral measurements such as inertial measurement units (IMUs) or eye trackers. We show examples of such simultaneous acquisition with MTw Awinda IMUs (Xsens, The Netherlands) (Fig. 5-A) and a Tobii Pro Fusion (Tobii, Sweden) eye tracker (Fig. 5-B).

**Figure 5.**
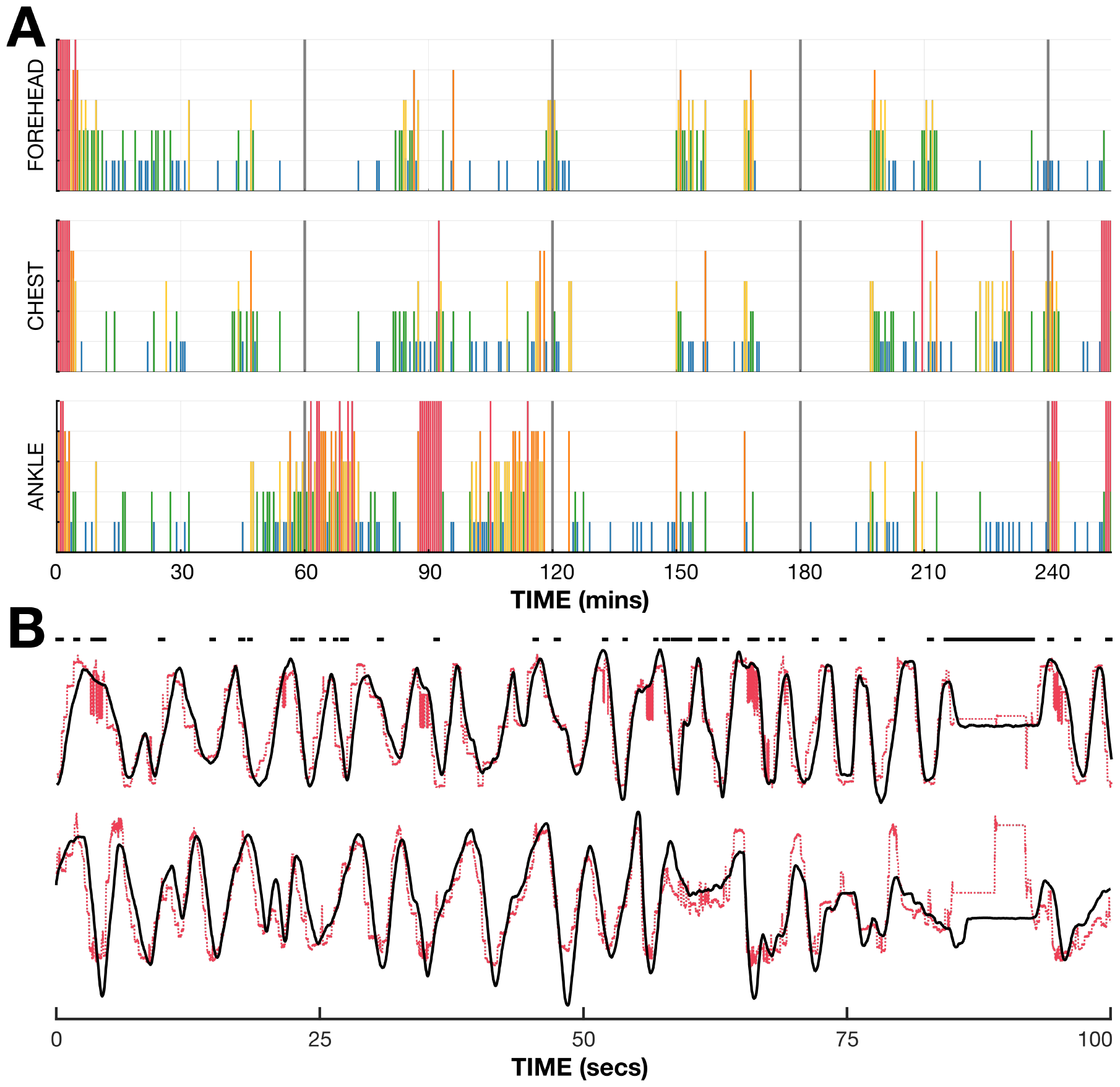
Synchronized acquisition of signals from different devices. A: Movements from one subject during approximately four hours of sleep. The top panel shows forehead movements detected using our device; and the center and bottom panels show concurrent chest and ankle movements detected using MTw IMUs, respectively. Each colored vertical line represents a 30-second time period, and the color and length of each line gives the magnitude of movement. As expected, movements detected at these three positions are similar but not identical. B: Head and eye movements from one subject who focused on different parts of a computer screen during a 100-second period. Black solid time courses give forehead movements detected using our device; red dashed time courses give eye movements detected using the Tobii Pro Fusion eye tracker. Top and bottom panels show horizontal and vertical movements, respectively. Black dots on top indicate the times during which the eye tracker could not detect eye movements (presumably due to eye blinks). Head movements detected by our device closely track the eye movements detected by the eye tracker.

### 3.3. Evoked Potentials Resulting from Visual Stimulation

The third context involves the capability to present auditory/visual stimuli, and to synchronize the timing of the stimuli with EEG data collection. This capability enables closed-loop neurotechnology applications (such as Doan et al. (2021); Musso et al. (2022)) that depend on evaluation of evoked potentials. Our platform can readily implement such protocols.

To showcase a typical example, we implemented a visual oddball paradigm that sequentially presented visual stimuli that were of one of two types. The first type of stimulus was a picture of a face (standard stimulus); the second type of stimulus was a picture of a zebra (oddball stimulus). Each stimulus was presented for 150 ms, and the inter-stimulus interval (ISI) randomly varied between 340-640 ms. The sequence of stimuli was block-randomized in blocks of 10. Each block contained a random sequence of 8 standard and 2 oddball stimuli. The subjects were asked to count the total number of oddball stimuli presented throughout the task.

Fig. 6 shows average evoked responses to oddball (red trace) and standard (blue) visual stimulation. Shaded areas indicate the standard error. As expected from the recording/reference locations on the forehead and consistent with the findings in Schalk et al. (2023), the responses are smaller in amplitude and visually different than those measured with more common montages (e.g., Cz referenced to the earlobe). At the same time, they clearly detect EEG responses to visual stimulation, and they are different for oddball and standard stimulation. This example demonstrates that our platform can readily implement such visual stimulation synchronized to wireless EEG acquisition, and our device with its forehead electrode montage can detect resulting EEG responses.

**Figure 6.**
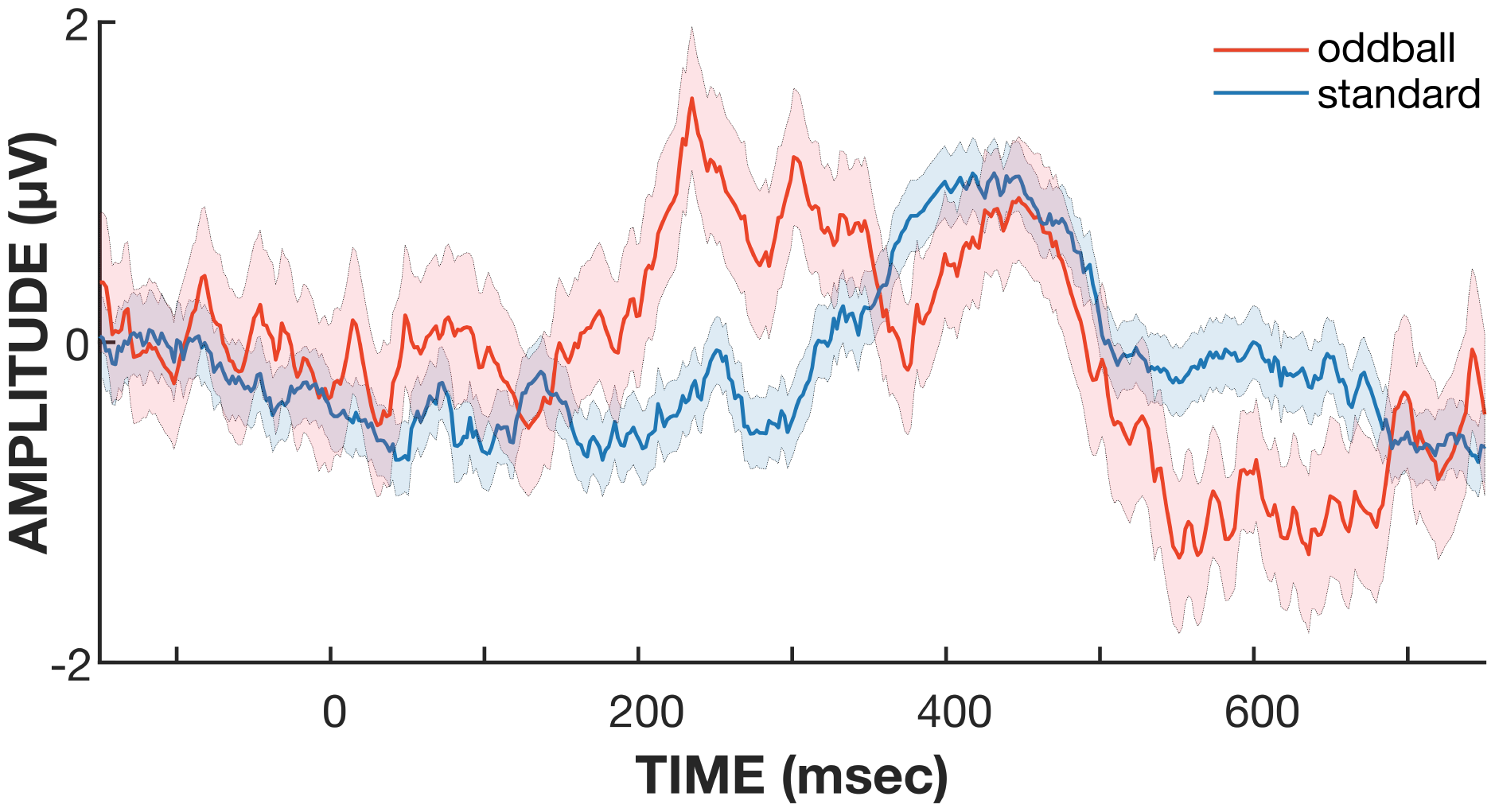
Evoked responses resulting from visual stimulation. We presented a series of oddball and standard visual stimuli (see text for details). Red and blue traces give the resulting average evoked responses, respectively, and shaded areas indicate the standard error. Visual stimulation generates clear evoked potentials, and those are different between the oddball and standard stimuli.

## 4. Discussion

### 4.1. Summary

In this paper, we describe our work toward the development of a general-purpose non-invasive neurotechnology research platform that is based on our multi-modal recording device, an associated API, and an interface to BCI2000 software. As the examples herein demonstrate, we implemented several protocols that demonstrate the platform’s capabilities and that will facilitate research into home-based sleep monitoring and modulation techniques as well as other neurotechnologies. Thus, our successful demonstration brings us closer to the day when it will be possible to more practically interact with the human brain with sophisticated recording and stimulation protocols.

### 4.2. Our Approach to Translation

It is undisputed that the identification and development of neurotechnologies for practical home use is a challenging enterprise. It requires the simultaneous optimization of neuroscientific protocol, hardware, software, and signal processing/AI algorithms, and the design of an easy-to-use solution that can deliver clear (and otherwise unobtainable) value to the user.

To date, the predominant approach to identifying and testing neurotechnology candidates has been to conduct laboratory studies using research equipment and highly trained personnel, thereby greatly sacrificing practicality in favor of system performance. The underlying logic is to describe the validity of a certain neurotechnology candidate first, and then to optimize the approach for home use later.

By proposing the platform described herein, we imply and promote an entirely different approach. We suggest to begin with a system that, while clearly limited in electrode coverage, is much more practical. With this increase in practicality, more neurotechnology candidates can be evaluated, and successful candidates have a more direct path to adoption.

We are fully aware that there may be efficacious applications of EEG technology that critically depend on trained personnel and/or complex EEG montages or other aspects of system configuration. In this case, it may be impossible to realize them with a more reduced system such as ours, or otherwise make them more practical. We argue that, if the primary goal is to produce neurotechnology solutions that eventually will improve the lives of many people, both efficacy and practicality are critical and need to be considered from the beginning.

### 4.3. Availability

We completed an initial complete prototype of our platform, and have begun to share it with a distinct set of collaborators. We anticipate to continue sharing it with select partners, and expand availability over time as the capabilities and robustness of our platform further increase. Please send inquiries about the availability of our platform to.

### 4.4. Conclusions and Outlook

Neurotechnologies have the potential to improve many people’s lives, but to date, we have barely scratched the surface of these opportunities. An important reason for this lack of widely accessible solutions is the lack of a capable hardware/software platform that makes it easy to develop and evaluate different non-invasive neuromodulation approaches. The work described in this paper illustrates our initial steps towards addressing this issue. At the same time, much work is left to be done. To maximize the potential impact of our work, we need to prepare our platform for more widespread dissemination and then make it widely available together with appropriate documentation and training.

## 5. Acknowledgements

We gratefully acknowledge funding from the Tianqiao and Chrissy Chen Institute (TCCI), the National Natural Science Foundation of China (82272116 (LC), 52105030 (GL)), the Shanghai Natural Science Foundation (23ZR1430900 (GL)), and the National Key Research and Development Program (2021YFC2501404 (HY)). We also thank Jason Reindorp (TCCI) and Tony Larsson (deDesigned) for their help with illustrations, and Duofu Liu (TCCI) for his support of the hardware project.

## References

Alkhoury L, Choi J w, Wang C, Rajasekar A, Acharya S, Mahoney S, Shender B S, Hrebien L and Kam M 2021 Journal of Clinical Monitoring and Computing 35(4), 797–813. URL: 10.1007/s10877-020-00539-2

Arnal P J, Thorey V, Debellemaniere E, Ballard M E, Bou Hernandez A, Guillot A, Jourde H, Harris M, Guillard M, Van Beers P, Chennaoui M and Sauvet F 2020 Sleep 43(11), zsaa097. URL: 10.1093/sleep/zsaa097

Beacon Biosignals 2023 ‘Dreem’. URL: https://beacon.bio/dreem-headband

Benjafield A V, Ayas N T, Eastwood P R, Heinzer R, Ip M S, Morrell M J, Nunez C M, Patel S R, Penzel T, Pépin J L et al. 2019 The Lancet Respiratory Medicine 7(8), 687–698.

Biswas D, Sim ões-Capela N, Van Hoof C and Van Helleputte N 2019 IEEE Sensors Journal 19(16), 6560–6570.

Brunner P and Schalk G 2018 in C. S Nam, A Nijholt and F Lotte, eds, ‘Brain-Computer Interfaces Handbook: Technological and Theoretical Advances’ CRC Press, Taylor & Francis Cambridge, MA, USA pp. 323–336.

Bundy D T, Souders L, Baranyai K, Leonard L, Schalk G, Coker R, Moran D W, Huskey T and Leuthardt E C 2017 Stroke 48(7), 1908–1915.

Compumedics 2023 ‘Somfit/Somfit Pro’. URL: https://www.compumedics.com.au/en/products/somfit/

da Silva Souto C F, Pätzold W, Wolf K I, Paul M, Matthiesen I, Bleichner M G and Debener S 2021 Frontiers in Digital Health 3, 688122.

De Fazio R, Mattei V, Al-Naami B, De Vittorio M and Visconti P 2022 Micromachines 13(8). URL: https://www.mdpi.com/2072-666X/13/8/1335

de Kock J and Tarassenko L 1991 Journal of Biomedical Engineering 13(1), 61–66. URL: https://www.sciencedirect.com/science/article/pii/014154259190046A

deAguiar Neto F S and Rosa J L G 2019 Neuroscience & Biobehavioral Reviews 105, 83–93.

Doan D N T, Ku B, Choi J, Oh M, Kim K, Cha W and Kim J U 2021 Frontiers in Aging Neuroscience 13, 180.

EMOTIV Inc. 2023 ‘Emotiv’. URL: https://www.emotiv.com/

FlectoThink 2022 ‘Airdream’. URL: https://www.flexolinkai.com/about.html

Garcia-Molina G, Tsoneva T, Jasko J, Steele B, Aquino A, Baher K, Pastoor S, Pfundtner S, Ostrowski L, Miller B et al. 2018 Journal of Neural Engineering 15(6), 066018.

g.tec 2023 ‘g.tec medical engineering’. URL: https://www.gtec.at/product/

Herff C, Heger D, De Pesters A, Telaar D, Brunner P, Schalk G and Schultz T 2015 Frontiers in Neuroscience 9, 217.

Hunan VentMed Medical Technology 2017 ‘Sf-c’. URL: https://www.ventmed.cn/product.aspx?ProductTypeId=1006

InteraXon Inc. 2023 ‘muse’. URL: https://choosemuse.com/

Jeong J 2004 Clinical Neurophysiology 115(7), 1490–1505.

Kwon S, Kim H S, Kwon K, Kim H, Kim Y S, Lee S H, Kwon Y T, Jeong J W, Trotti L M, Duarte A et al. 2023 Science Advances 9(21), eadg9671.

Kwon S, Kim H and Yeo W H 2021 Iscience 24(5).

Leuthardt E, Schalk G JR J W, Ojemann J and Moran D 2004 1(2), 63–71.

Madgwick S O H 2010. URL: https://api.semanticscholar.org/CorpusID:2976407

Mahony R, Hamel T and Pflimlin J M 2008 IEEE Transactions on Automatic Control 53(5), 1203–1218.

Miller K J, Schalk G, Fetz E E, Den Nijs M, Ojemann J G and Rao R P 2010 Proceedings of the National Academy of Sciences 107(9), 4430–4435.

Morin C M and Jarrin D C 2022 Sleep Medicine Clinics 17(2), 173–191.

Musso M, Hübner D, Schwarzkopf S, Bernodusson M, LeVan P, Weiller C and Tangermann M 2022 Brain Communications 4(1), fcac008.

Nakamura T, Goverdovsky V, Morrell M J and Mandic D P 2017 IEEE Journal of Translational Engineering in Health and Medicine 5, 1–8.

Neurable 2023 ‘neurable’. URL: https://www.neurable.com/

NeuroSky Inc. 2023 ‘neurosky’. URL: http://neurosky.com/

Ngo H V V, Martinetz T, Born J and Mölle M 2013 Neuron 78(3), 545–553.

OpenBCI 2022 ‘openbci’. URL: https://openbci.com/

Pankaj, Kumar A, Komaragiri R and Kumar M 2022 Archives of Computational Methods in Engineering 29(2), 921–940. URL: 10.1007/s11831-021-09597-4

Papalambros N A, Weintraub S, Chen T, Grimaldi D, Santostasi G, Paller K A, Zee P C and Malkani R G 2019 Annals of Clinical and Translational neurology 6(7), 1191–1201.

Rundo J V and Downey III R 2019 Handbook of Clinical Neurology 160, 381–392.

Santostasi G, Malkani R, Riedner B, Bellesi M, Tononi G, Paller K A and Zee P C 2016 Journal of Neuroscience Methods 259, 101–114.

Schalk G, McFarland D, Hinterberger T, Birbaumer N and Wolpaw J 2004 IEEE Transactions on Biomedical Engineering 51(6), 1034–1043.

Schalk G and Mellinger J 2010 A Practical Guide to Brain-Computer Interfacing with BCI2000 1st edn Springer London, UK.

Schalk G, Shao S, Xiao K and Wu Z 2023 Biomedical Physics & Engineering Express .

Senaratna C V, Perret J L, Lodge C J, Lowe A J, Campbell B E, Matheson M C, Hamilton G S and Dharmage S C 2017 Sleep Medicine Reviews 34, 70–81.

Shenzhen EEGSmart Technology 2019 ‘UmindSleep’. URL: http://www.eegsmart.com/en/UMindSleep.html

Surani S and Taweesedt P 2022 ‘Obstructive sleep apnea: New perspective’.

Tabar Y R, Mikkelsen K B, Shenton N, Kappel S L, Bertelsen A R, Nikbakht R, Toft H O, Henriksen C H, Hemmsen M C, Rank M L et al. 2023 Frontiers in Neuroscience 17, 987578.

Wearable Sensing 2023 ‘Wearable Sensing’. URL: https://wearablesensing.com/

Wilson J A, Mellinger J, Schalk G and Williams J 2010 IEEE Transactions on Biomedical Engineering 57(7), 1785–1797.

Wolpaw J and McFarland D 2004 Proc Natl Acad Sci U S A 101(51), 17849–17854.

Wolpaw J and Wolpaw E, eds 2012 Brain-Computer Interfaces: Principles and Practice Oxford University Press New York.

Wukitsch M W, Petterson M T, Tobler D R and Pologe J A 1988 Journal of Clinical Monitoring 4(4), 290–301. URL: 10.1007/BF01617328

Xu Y, De la Paz E, Paul A, Mahato K, Sempionatto J R, Tostado N, Lee M, Hota G, Lin M, Uppal A et al. 2023 Nature Biomedical Engineering pp. 1–14.

Ye Y, Cheng Y, He W, Hou M and Zhang Z 2016 IEEE Sensors Journal 16(19), 7133–7141.

